# Filopodia-like Structures are Essential for Steroid Release

**DOI:** 10.1101/2024.10.10.617560

**Authors:** Eléanor Simon, Raphaël Bonche, Yassine Maarouf, Marie-Paule Nawrot-Esposito, Nuria Magdalena Romero

**Author notes:** Contributed equally (Eléanor Simon, Raphaël Bonche). Correspondence should be addressed to N.R.

## Abstract

Steroid hormones, crucial for development and physiology, were traditionally believed to diffuse passively through membranes. However, recent evidence shows insect steroid ecdysone being secreted via regulated exocytosis, but the mechanisms ensuring successful hormone release into circulation remain unclear. Our study identifies specialized membrane protrusions, filopodia-like structures named “hormonemes,” in the Drosophila prothoracic gland as essential for steroid vesicle release. Confocal imaging reveals that these actin- and tubulin-rich structures form a membrane-intertwined basal domain critical for secretion. Disrupting filopodia by interfering with BM interactions—perlecan or β-integrin—or filopodia-specific protein expression—α-actinin—significantly reduces ecdysone signaling by impairing its release, despite proper production in the gland. Additionally, filopodia dynamics, such as length and density, align with secretion timing and hormone circulating levels, suggesting their role in synchronizing release with physiological needs. The systematic presence of membrane protrusions in steroid-secreting glands across species prompts a comprehensive re-evaluation of steroid release mechanisms.

## Introduction

Steroid hormones are essential bioactive molecules that regulate a wide range of physiological processes in both animals and plants, including immune response, salt and water balance, glucose metabolism, and developmental sexual maturation^1,2^. Rapid fluctuations in circulating steroid hormone levels are essential for exerting their effects on distant target tissues. Despite their critical roles, driving steroid hormone release from endocrine tissues remains poorly understood. Endocrine gland cells are typically organized in cords or clumps, attached laterally to neighboring cells, and often lack classical apical-basal polarity. This unique organization raises an important question: How do endocrine glands efficiently direct rapid surges of hormonal secretion outward? While it was traditionally thought that steroid hormones diffuse across lipid bilayers, recent studies have revealed a more complex and regulated process. For instance, in *Drosophila*, the release of the steroid hormone Ecdysone involves a regulated vesicular trafficking mechanism dependent on calcium signaling, Rab3, synaptotagmin 1 (Syt1), and an ABC transporter, Atet, which facilitates Ecdysone transport across the lipid bilayer^3^. In humans, steroid hormones are detected in blood and urine within exovesicles^4^. These findings suggest that steroid hormone secretion is regulated, potentially involving multiple pathways depending on the tissue or signaling context.

Regulated exocytosis and multivesicular body (MVB) recycling pathways require a fusion event between the plasma membrane and the vesicle or MVB membrane, which poses a significant energetic challenge for cells^5^. The mass secretion of steroids during maturation exacerbates this issue, highlighting the need for efficient mechanisms to reduce energy demands. Cells can mitigate these demands by altering their lipid composition^6,7^, membrane forces^8^, or curvature^9^, the energy required for stalk and pore formation during membrane fusion is lower at higher curvatures membranes^9,10^. Therefore, by optimizing these critical membrane properties, cells might enhance membrane fusion efficiency, facilitating vesicle secretion. Interestingly, electron microscopy studies from the 1960s on the crab endocrine Y-organ revealed cell surface “irregularities” with steroid-containing vesicles^11^, suggesting a direct link between these membrane structures and steroid hormone secretion. Similar observations have been made in the prothoracic gland of lepidopterans^12^ and dipterans^13^, as well as in vertebrate steroid-secretory glands like the adrenal cortex^14^ and luteal cells^15–17^. These irregularities, described as blebs, microvilli, or filopodia depending on the tissue and fixation technique (note that membrane projection properties are affected by fixation methods^18^, appear to be common features related to steroid secretion across different species. This crucial aspect has been completely overlooked.

Filopodia, cytonemes, microvilli, and blebs are dynamic membrane protrusions that play essential roles in cellular processes, including migration, cell adhesion, cell-to-cell communication, signaling, and wound healing ^19,20^. Primarily composed of tightly bundled actin filaments, these slender, finger-like structures can also incorporate microtubules. The interplay between actin and microtubules is vital for maintaining their stability^21,22^. Notably, microtubules serve as scaffolds for vesicle transport within protrusions, raising the question of whether filopodia containing microtubules not only direct the transport of steroid vesicles outward but also provide local plasma membrane curvature^23^ necessary for membrane fusion and, consequently, vesicle secretion.

In this study, we explore the role of membrane projections in mediating the secretion of Ecdysone in the Drosophila prothoracic gland (PG). We provide a detailed description of the PG’s structure, highlighting a bilayer organization surrounded by a basement membrane (BM) lacking classical apical-basal polarity. Through clonal analyses and *ex vivo* experiments, we identified filopodia-like membrane projections concentrated in the sub-BM region. These filopodia contain actin, β-integrin, and microtubules, forming an asymmetric membrane network. Disruption of the endocytic/MVB pathway via Rab11 down-regulation altered PG cell morphology, reduced filopodia presence, and delayed development. Furthermore, the asymmetric distribution of filopodia depends on BM components like Trol and β-integrin. Filopodia size and density correlate with Ecdysone secretion peaks, with significant structural changes during the late larval stages. We also found that filopodia contain Ecdysone secretory machinery, including Syt1 and Atet. Silencing filopodia-specific components resulted in developmental delays and reduced Ecdysone secretion, confirming their crucial role in secretion. Filopodia dynamic significantly changes during the late larval stages regarding its size and density, which correlate with Ecdysone secretion peaks. In conclusion, our findings demonstrate that a specialized asymmetric membrane domain, formed by filopodia-like projections, is required for steroid hormone secretion in the insect prothoracic endocrine glands. This membrane network structure is essential for efficiently releasing ecdysone.

## Results

### The PG lacks classical apical-basal polarity but exhibits polarized secretion properties

To assess whether the PG shares traits with other endocrine glands in vertebrates and invertebrates, we analyzed the arrangement of PG cells in the ring gland using confocal microscopy. Z-stacks and orthogonal XZ/YZ sections revealed a well-organized tissue with two layers: a dorsal layer containing a more significant number of smaller cells and a ventral one (**Figure 1A, Supp Figure 1A**). Only the ventral layer is innervated by tracheal cells (**Figure 1A**). A 3D visualization is show in **Figure 1A’**. As expected, no lumen was present between the layers (**Supp Figure 1B-C**). PTTH neurons were restricted to the interface of the two layers (**Figure 1B,B’,B’’**), and the PTTH receptor, Torso, was nicely concentrated at postsynaptic regions of the middle cell membrane (**Figure 1C**). Using the *terribly reduced optic lobes* (*trol*)-GFP knock-in reporter (a homolog of vertebrate Perlecan), we observed a BM surrounding the gland (**Figure 1D, D’**), also containing Viking (Vkg), Nidogen (Ndg) and Laminin-B1 (LanB1) confirmed by knocking GFP fusion protein reporters (**Supp Figure 1D-D’’’**). Unexpectedly, we found BM protein dots within the tissue along the lateral and middle PG cell membranes (**Figure 1D, D’** arrowheads, **Supp Figure 1D-D’’’**). These dots are Collagen IV intercellular concentrations (CIVIC) structures since a colocalization is observed between Cg25C and trol-GFP, vkg-GFP, Ndg-GFP and LanB1-GFP (**Supp Figure 1D-D’’’**). Despite its epithelial origin, PG cells exhibit CIVIC structures which is typically associated with unpolarized mesodermal fat body cells ^24^. Next, we examined whether PG cells display apical-basal polarity by staining them for epithelial markers. Apical markers bazooka (baz)-GFP and crumbs (crb)-YFP were absent from the membranes (**Supp Figure 1E-F**), and no crb expression was detected using an anti-crb antibody (not shown). Adherens junction proteins Armadillo (Arm), E-cadherin (Ecad), the septate junction markers Coracle (Cora) and Disc large 1 (Dlg1), as well as the βPS integrin subunit encoded by *myospheroid* (*mys*) were uniformly distributed along the membrane (**Figure 1E**). The N-cadherin (CadN) was only expressed in the CA (**Supp Figure 1G**). These findings strongly suggest that *Drosophila* PG lacks typical apical-basal polarity and preserves mesodermic properties acquired at the late embryonic stages^25^, a characteristic also seen in other endocrine glands like the adrenal gland ^26^. Interestingly, a Z-stack analysis of PG cells overexpressing Syt1-GFP revealed that Syt1-GFP is concentrated twice as much in the sub-BM region compared to the lateral or midline membranes (**Figure 1F**), indicating a polarized secretion. Therefore, three distinctive and polarized cell membrane domains emerge at the PG: a sub-BM membrane domain right below the BM enriched of Syt1 secretory marker, a lateral membrane one only in contact laterally with the neighboring cells of the same layer, and the middle membrane in contact with the PTTH neuron projections and cells of the other layer (schematic in **Figure 1G**).

**Figure 1.**
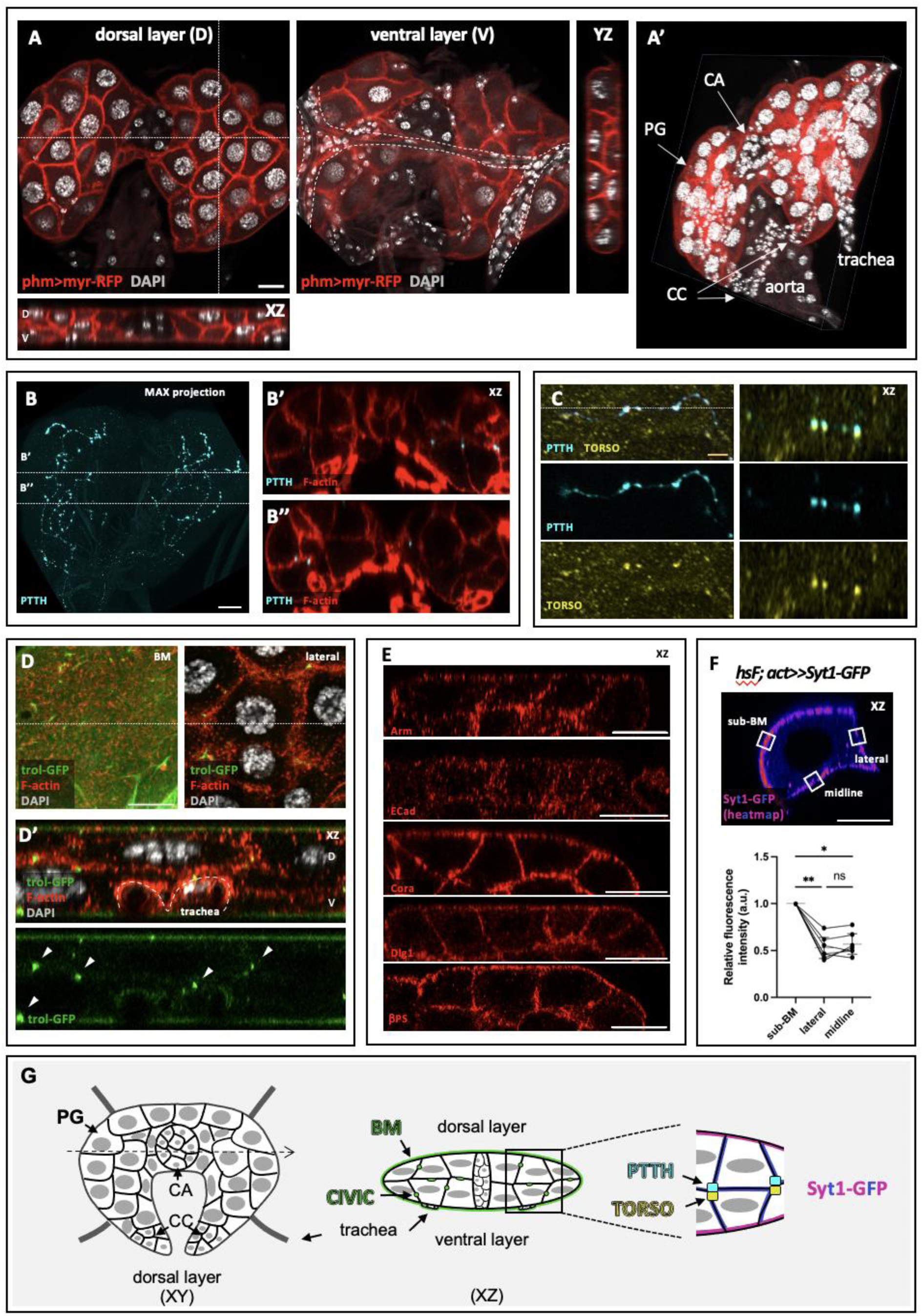
The PG is a two-cell layers organ surrounded by a BM. **(A,A’)** The PG is organized into a dorsal and ventral layer (A). 3D visualization of the ring gland (A’) composed of the PG (prothoracic gland), the corpora allata (CA) and the corpora cardiaca (CC). **(B-B’’)** Maximal projection of prothoracicotropic hormone (PTTH) staining (B) and two Z-sections revealing that PTTH projects its axons between the midline of the PG (B’, B’’). **(C)** Synaptic PTTH buttons reveal PTTH in the pre-synapse and Torso in the post-synapse. **(D, D’)** Expression of *trol-GFP* in the BM plan and in a lateral plan (D). The XZ section (D’) facilitates the observation of the *trol-GFP* dots. Those dots are CIVICs. **(E)** Z-sections reveal that the adherens junction markers Arm and ECad are located all around the PG cells. The septate junction markers and Dlg1 and the Cora and the βPS are also homogenously distributed in the PG cells. **(F)** Heatmap of Syt1-GFP in an XZ section of a cell revealed that the relative fluorescence intensity is higher in the sub-BM region compared to the lateral or midline membrane. The quantification is shown below. Scale bar: 20 μm. **(G)** Schematic representation of a PG dorsal layer and XZ cut.

### PG cells exhibit polarized features highlighted by an uneven distribution of filopodia in the sub-BM domain

We induced random fluorescent clones within the PG using the flip-out technique and visualized individual cell shapes. Confocal images revealed that PG cells exhibit yet unidentified membrane extensions of several µm resembling filopodia (**Figure 2A**). These fluorescence extensions are visible only when interfacing with a neighboring “black” cell that is non-expressing the fluorescent signal. These projections were predominantly observed in the sub-BM region, where PG cells face the BM, and were largely absent from lateral and middle membranes. This was evidenced by the distinct distribution observed using two independent membrane markers such as myristoylated (myr)-GFP (**Figure 2B**) and the PH domain of Phospholipase C (PLC)-GFP (**Figure 2C**). Overexpression of myr-GFP and PLC-GFP in the fat body, another highly secretory larval tissue composed of polyploid cells, resulted in smooth edges without the presence of filopodia-like (**Figure 2D**), indicating that these membrane projections are specific to PG cells. To further explore the organization of these membrane projections, we used the CoinFlp combined with GRASP (GFP Reconstitution Across Synaptic Partner) systems^27,28^, allowing us to observe interactions between neighboring PG cells. When complementary parts of GFP (spGFP^1-10^ and sp-GFP^1^) were expressed in adjacent cells, GFP reconstitution revealed intertwined membrane projections primarily located in the sub-BM region (**Figure 2E**). In contrast, in lateral sections, a distinct stripe of GFP signal is evident, reflecting the reduced number of those “projections” in this plane, confirming a clear asymmetry in the projection distribution between cells.

**Figure 2.**
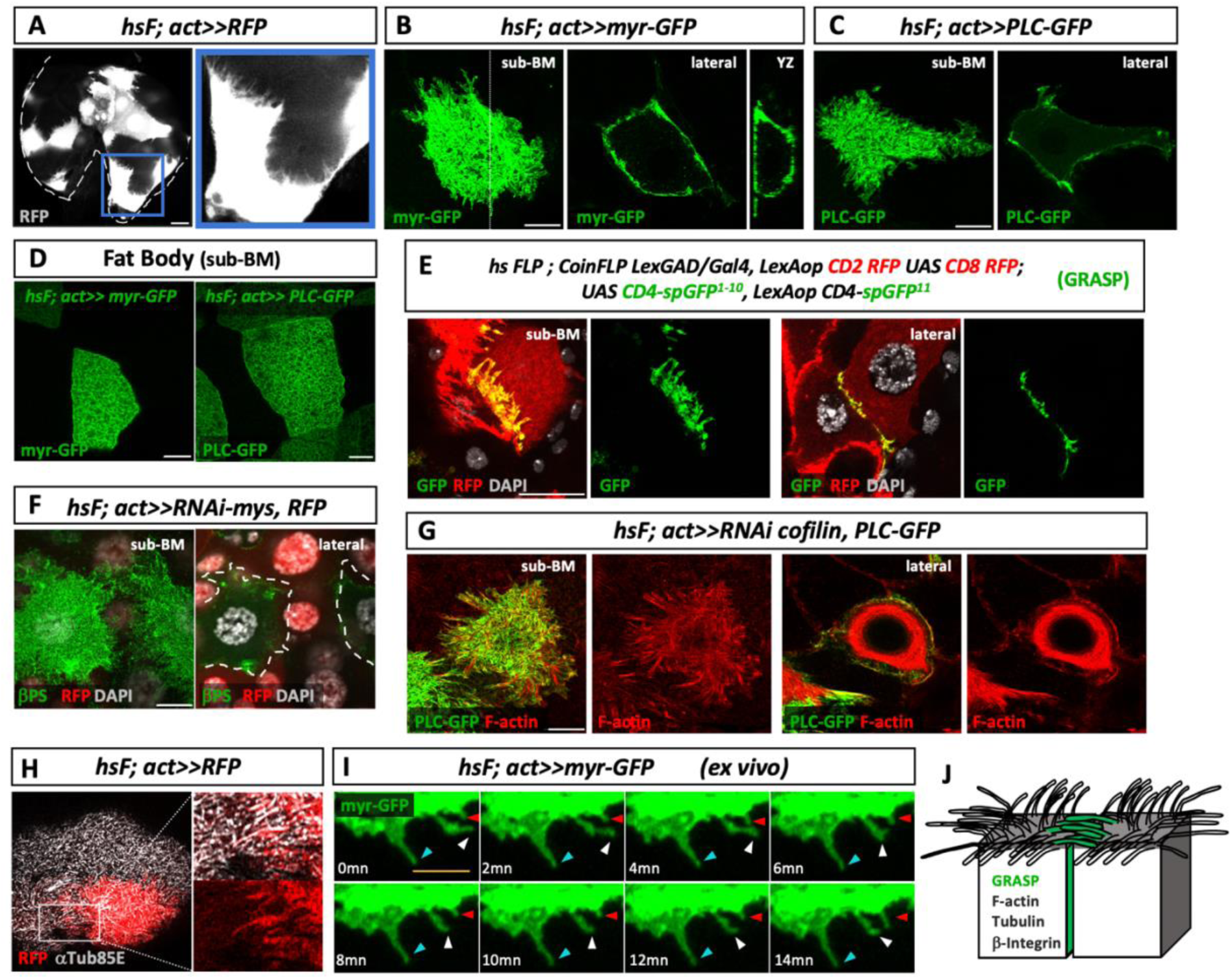
PG Cells Emit Actin- and Tubulin-Based Filopodia. **(A)** Flip-out clones randomly expressing the fluorescent marker RFP in the PG Cells emit membrane extensions only visualized when neighboring cells do not express the RFP (see insert). The picture corresponds to a maximal projection of a Z-stack. **(B)** Flip-out clone expressing the membrane marker myr-GFP. Long filopodia are observed on the sub-BM side, while fewer and shorter filopodia are observed in a lateral plane. The XZ section reveals a long filopodium present in the sub-BM region. The lateral side of the cell nor the side facing the midline emit obvious filopodia. **(C)** The flip-out clone expresses the membrane marker PLC-GFP, which shows mainly filopodia in the sub-BM region. **(D)** Flip-out clones induced in the fat body marked either by myr-GFP (left) or PLC-GFP (right). In the sub-BM plane, no filopodia are observed. **(E)** Two neighboring cells (marked positively with the RFP thanks to the CoinFLP system) expressing each a part of the GFP (spGFP^1-10^ or sp-GFP^11^; GRASP technique). On the sub-BM side, reconstructed GFP revealed physical integrations between filopodia of the two neighboring cells. In contrast, a stripe of GFP signal corresponding to a minimal physical interaction between the two cells is observed on the lateral plane. **(F)** Flip-out clones are marked positively by the RFP and express an RNAi against *mys*. The PG is stained against the βPS (green). Endogenous filopodia decorated by βPS are observed in the two WT cells (RFP negative) in the sub-BM plan. In the lateral plane, βPS is restricted to the cortex. **(G)** Flip-out clone was marked positively by PLC-GFP and expressed an RNAi against cofilin, impeding the destabilization of F-actin. In the sub-BM plane, long filopodia containing F-actin are observed. In the lateral plane, no actin-based filopodia are observed despite F-actin stabilization. **(H)** A flip-out RFP clone showing a microtubule presents in the filopodia (inserts). **(I)** *ex vivo* experiment showing filopodia tracking by expressing myr-GFP. The blue, white, and red arrows point out the stability of the filopodia through time. White scale bar: 20 μm; yellow scale bar: 5 μm. **(J)** Schematic representation of the filopodia distribution in PG cells contains F-actin, tubulin, and βPS.

These membrane projections resemble filopodia, thin, finger-like structures formed by bundles of actin filaments which interact with the ECM through the transmembrane protein βPS. To determine the membrane projection are indeed filopodia, we aim to visualize the localization of βPS. To do so, we induced random clones silencing *mys*, the gene encoding for βPS, in PG cells to employ the “black” background clonal method to visualize filopodia. Under this mosaic expression pattern for βPS, we reveal the presence of endogenous filopodia decorated by βPS in the sub-BM region (**Figure 2F**).

After, we proceeded to analyze the presence of F-actin using phalloidin staining. Unexpectedly, actin filaments were not visible in the sub-BM region; instead, we observed a diffuse punctate pattern, even though actin was clearly present in the cell cortex (data not shown). We suspected that a balance between F-actin polymerization and depolymerization might have hindered its detection. To test this, we reduced actin dynamics by creating random prothoracic gland cell clones with silenced cofilin (twinstar), a protein responsible for F-actin depolymerization^29^. Under these conditions, we observed a significant accumulation of F-actin along the sub-BM membrane projections (**Figure 2G**).^18,30,21,22^

Although filopodia typically contain only actin and βPS, some exceptions with microtubules have been reported^18,30^. Given the high expression of αTub85E in active secretory PGs ^31^, we stained the gland for this α-tubulin subunit. Surprisingly, we detected microtubules within the filopodia-like structures (**Figure 2H**). Moreover, a lateral section revealed the slight presence of αTub85E in the cytoplasm in a uniform, non-polymerized manner, suggesting the absence of αTub85E microtubules within the cytoplasm (**Supp Fig 2A and B**). Since filopodia are delicate and sometimes do not survive fixation well, we performed *ex vivo* experiments to visualize their real features and dynamics. Time-lapse imaging over 15 minutes confirmed the presence of numerous filopodia and revealed their low motility (**Figure 2I and Movie 1**). This aligns with previous findings showing that the association between F-actin and microtubules provides mechanical stability ^21,22^. Our findings demonstrate that PG cells possess filopodia-like extensions composed of F-actin, integrins, and microtubules, which exhibit polarized distribution predominantly below the BM. This organization indicates that PG cells possess polarized properties based on an asymmetric distribution of cytoskeleton components (**Figure 2J**).

### The MVB/recycling pathway is required for the asymmetric PG cells membrane identity and for timing development

Previous electron microscopy studies have revealed an uneven distribution of multivesicular bodies (MVBs) primarily near the cell cortex on the BM side of the PG cells^13^. Since MVBs are visualized only in highly secretory PG cells^13^ we evaluated the function of the PG MVB/recycling pathway on timing development, which is defined by the Ecdysone surges. Our experiments revealed that down-regulation of Rab11 leads to larval arrest or an 80-hour delay in development (**Figure 3A**). Additional experiments silencing other MVB pathway components, including Vps25, Vps36, and Shrb, confirmed a significant developmental delay when this pathway is affected (**Figure 3B**). These findings underscore the critical role of the MVB pathway in the PG functionality; therefore, we investigated potential mechanisms behind these effects. First, we evaluate the direct secretion of ecdysone into exovesicles, by altering the MVB/recycling pathway. Indeed, human steroid hormones have been detected in exosomes, whose secretion depends on the MVB pathway^4^. Therefore, we expressed the small GTPase Rab11-GFP in PG cells and purified circulating exosomes from the hemolymph using an ultracentrifugation protocol^32^. Even though we detected the presence of GFP-Rab11 PG containing exosomes in our exosome purified fraction by Western blot, our ecdysone measurement showed only negligible amounts of ecdysone in this fraction (**Supp Figure 3A**), indicating that the MVB/recycling pathway does not directly secrete ecdysone. This aligns with previously reported Rab3-dependent ecdysone secretory pathway ^3^.

**Figure 3.**
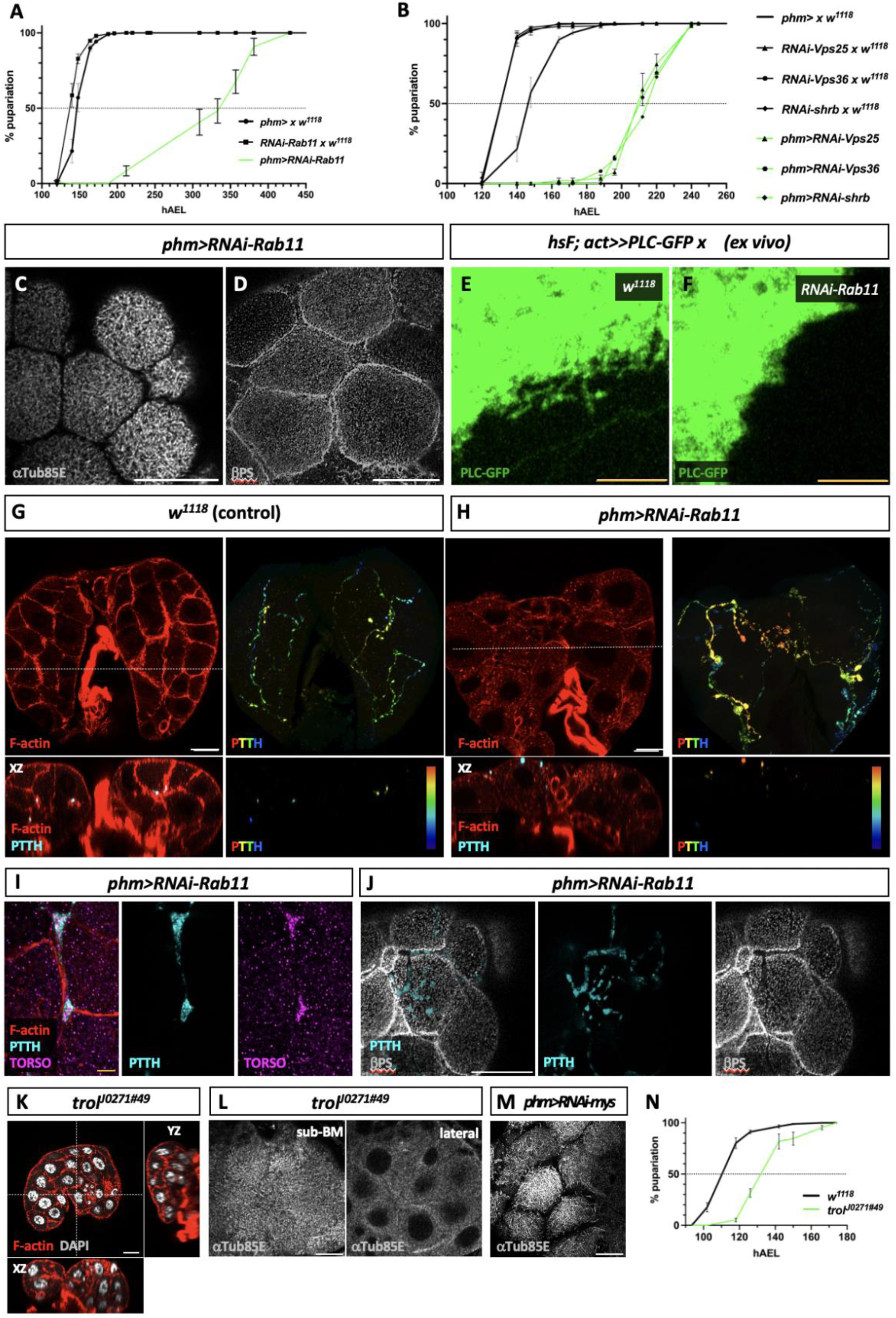
The role of the MVB pathway, integrins, and BM components in establishing asymmetric membrane identity and filopodia organization in PG cells. **(A)** Inhibition of Rab11 results in an 80-hour delay in entering metamorphosis compared to the control. **(B)** Silencing of Vps25, Vps36, and shrub in PG cells resulted in a several-hour delay in entering metamorphosis compared to the control. **(C, D)** Images of *phm>RNAi-Rab11* PG displaying a lack of microtubules (C) and βPS filopodia extension (D) in the sub-BM region. (E, F) *ex vivo* PG In a control situation, filopodia are visible when the cell expresses PLC-GFP (E). However, they are not observed anymore when *Rab11* expression is down-regulated (F). **(G, H)** Comparison of control *w^1118^* PG (G) and *phm>RNAi-Rab11* PG (H) showing F-actin (single section) and PTTH staining (maximum projection of a Z-stack) processed with a depth color code from a Fiji plugin. The YZ cuts allow us to see the position of the PTTH axons between the PG bilayer in the control PG (G), while when the recycling pathway is altered, PTTH axons are also observed below the BM (H). **(I)** Axon from a *phm>RNAi-Rab11* PG at the sub-BM region. The PTTH staining co-localized with its receptor Torso, suggesting that these mislocalized synaptic buttons are functional. **(J)** A *phm>RNAi-Rab11* PG at the sub-BM region shows the mislocalized synaptic button containing PTTH that represses βPS expression. **(K)** Image of a *trol^JO271#49^* mutant PG Z sections reveals the loss of bilayer organization. **(L)** αTub85E staining in a *trol^JO271#49^*mutant PG shows short microtubules in the sub-BM region. In the lateral plan, αTub85E staining looks normal. **(M)** Down-regulation of *mys* impedes microtubule extension. **(N)** Developmental timing of *trol^JO271#49^* mutant vs *w^1118^*. The *trol^JO271#49^* mutant enters pupariation 24 hours after the control line. White scale bar: 20 μm; yellow scale bar: 5 μm.

Interestingly, the MVB/recycling pathway is also involved in the polarized distribution of adhesion proteins (reviewed in ^33,34^) and filopodia formation^35^. Therefore, we investigate whether this pathway impacts filopodia’s asymmetric and polarized distribution in Drosophila PG cells. We specifically manipulated the expression of Rab11 in the PG cells. We observed the formation of rounded cells lacking αTub85E staining between them (**Figure 3C**), suggesting a marked disruption of the filopodia network within the sub-BM region. Furthermore, reduced Rab11 levels in the PG resulted in well-defined cell borders and βPS accumulation at the cell edges (**Figure 3D**), suggesting a substantial reduction or complete absence of filopodia extensions. To confirm these findings and knowing that filopodia could be disrupted by fixation mechanism, we performed *ex vivo* analyses of PG cells, showing that, unlike the control group (**Figure 3E**), clone cells with Rab11 silencing for only 24 hours failed to extend filopodia as reported with the PLC-GFP-marked (**Figure 3F**). Consistent with the loss of identity of the PG cells membrane facing the BM, we also observed alterations at the PG midline when silencing Rab11. In a control situation, PTTH neurons project their axons between the midline (**Figure 3G**). However, down-regulation of Rab11 expression in the PG cells causes aberrant PTTH axon organization, often leading to mis-localized axons situated below the BM (**Figure 3H**). Remarkably, despite the mis-localization, PTTH staining, found in the pre-synapse, co-localizes with its receptor Torso in the post-synapse, suggesting the potential functionality of these mislocalized synaptic buttons (**Figure 3I**). It is interesting to note that when the PTTH neuron’s synaptic buttons are below the BM, we observed the absence of βPS expression at these membrane sites (**Figure 3J**), suggesting that filopodia extensions and PTTH neuron’s synaptic buttons are mutually exclusive structures. These findings underscore the critical role of the MVB pathway in the PG functionality, including its 3D organization, such as the uneven distribution of filopodia in the BM membrane domain and the restricted interaction of the middle membrane domain with PTTH neurons; however, since Rab11 protein display several pleiotropic functions, we cannot directly link the presence of filopodia and the strong developmental timing phenotype observed. A hypothesis that we will further explore in the following approaches.

### The asymmetric distribution of filopodia in the sub-BM membrane of PG cells depends on integrins and Perlecan/Trol BM components

Given Rab11’s role in regulating βPS composition in the plasma membrane via the MVB/recycling pathway^36^, and considering that β-integrin interacts with BM components at filopodia^20^, we investigate whether this interaction influences the asymmetric distribution of filopodia in PG cells. To develop a better methodological approach to address this question, we first determine the source of *Drosophila* Perlecan, named Trol, in the BM and CIVIC structures. We conducted *trol-RNAi* clone expression analyses in the PG, marked with RFP, and observed that the presence of Trol in the internal dots of “CIVICs” disappeared, which are present in the unique WT non-RFP clone. However, in the same *trol-RNAi* clone analyses, we noticed that the levels of Trol at the BM were slightly higher autonomously in this WT cell (**Supp Figure 3B**). This contribution of the PG on Trol in the BM was further confirmed when silencing *trol* in all the PG using the RNAi technique. In this case, we observed a strong reduction of Trol in the CIVIC structures, whereas the one present in the BM is still evident but slightly reduced when compared with the control (**Supp Figure 3C**). Please note that Trol staining of control and *trol-RNAi* loss of function PGs was performed in the same tube for better comparison. Conversely, the *trol^JO27#49^* hypomorphic mutant shows a strong reduction in Trol expression at the BM, maintaining some signal at the internal CIVIC dots (**Supp Figure 3C**). These results indicate that the Trol protein present in the CIVICs is produced exclusively by the PG cells. In contrast, Trol in the BM comes partially from the hemolymph (as previously suggested in ^37^) with a non-depreciable autonomous PG contribution. Therefore, to address whether the BM plays a role in the asymmetric distribution of filopodia in the PG cells, we analyzed the PG in the hypomorphic allele *trol^JO27#49^*, which shows the absence of Trol in the BM (**Supp Figure 3C**). These PGs are relatively round compared to WT glands and show aberration in the Z plan. Indeed, three layers of cells replace the PG two-layer organization (**Figure 3K**). Examination of αTub85E staining in the sub-BM plan revealed short microtubules instead of the normal microtubes elongated observed in the controls (**Figure 2H**). In contrast, in the cytoplasm, the αTub85E pattern is normal (**Figure 3L**), suggesting a specific effect of Trol at the BM in αTub85E polymerization at the sub-BM plan. Accordingly, silencing *trol* specifically in the PG, affecting mainly the presence of Trol in the CIVICs, does not affect PG cell layers or αTub85E staining at the sub-BM (**Supp Figure 3D and E** respectively). Moreover, direct down-regulation of βPS expression in the PG cells causes a similar change in αTub85E staining in the sub-BM, shown by the reduced αTub85E signal at the PG cells boundaries (**Figure 3M**), suggesting the absence of αTub85E polymerization at the filopodia. Moreover, *trol^JO27#49^* mutants, as well as larvae that have βPS silenced exclusively in the PG cells, enter metamorphosis with an approximately 24-hour delay compared to control (**Figure 3N and Supp Figure 3F**), suggesting that alteration in the 3D organization in these glands is relevant for the establishment of the large Ecdysone peak inducing the onset of metamorphosis.

### PG filopodia contains the machinery for Ecdysone secretion

To investigate whether filopodia directly affects Ecdysone secretion, we analyzed Syt1 and Atet Ecdysone positive secretory vesicle markers in the filopodia^3^. We found that in PG clones expressing Syt1-GFP, the sub-BM membrane is enriched in filopodia decorated by Syt1-GFP, whereas in a lateral plan, we observe Syt1-GFP thin enrichment at the membrane. No filopodia (or few) are observed in the lateral membrane nor the middle one facing the midline (**Figure 4A and Movie 2**). A zoom into the sub-BM region enables us to appreciate the presence of Syt1 at the membrane of the filopodia and also in dots-vesicles along this filopodia (**Figure 4B**). Moreover, we overexpressed the fluorescent construct Ypet-Atet and observed the presence of Ypet-Atet fluorescent marker at the filopodia (**Figure 4C**) similarly to Syt1-GFP. To further confirm the presence of Syt1-GFP vesicles in filopodia, we performed *ex vivo* experiments, where we nicely observed the directional movement of those vesicles through filopodia (**Figure 4E**). The findings suggest that filopodia contains the necessary components to secrete Ecdysone.

**Figure 4.**
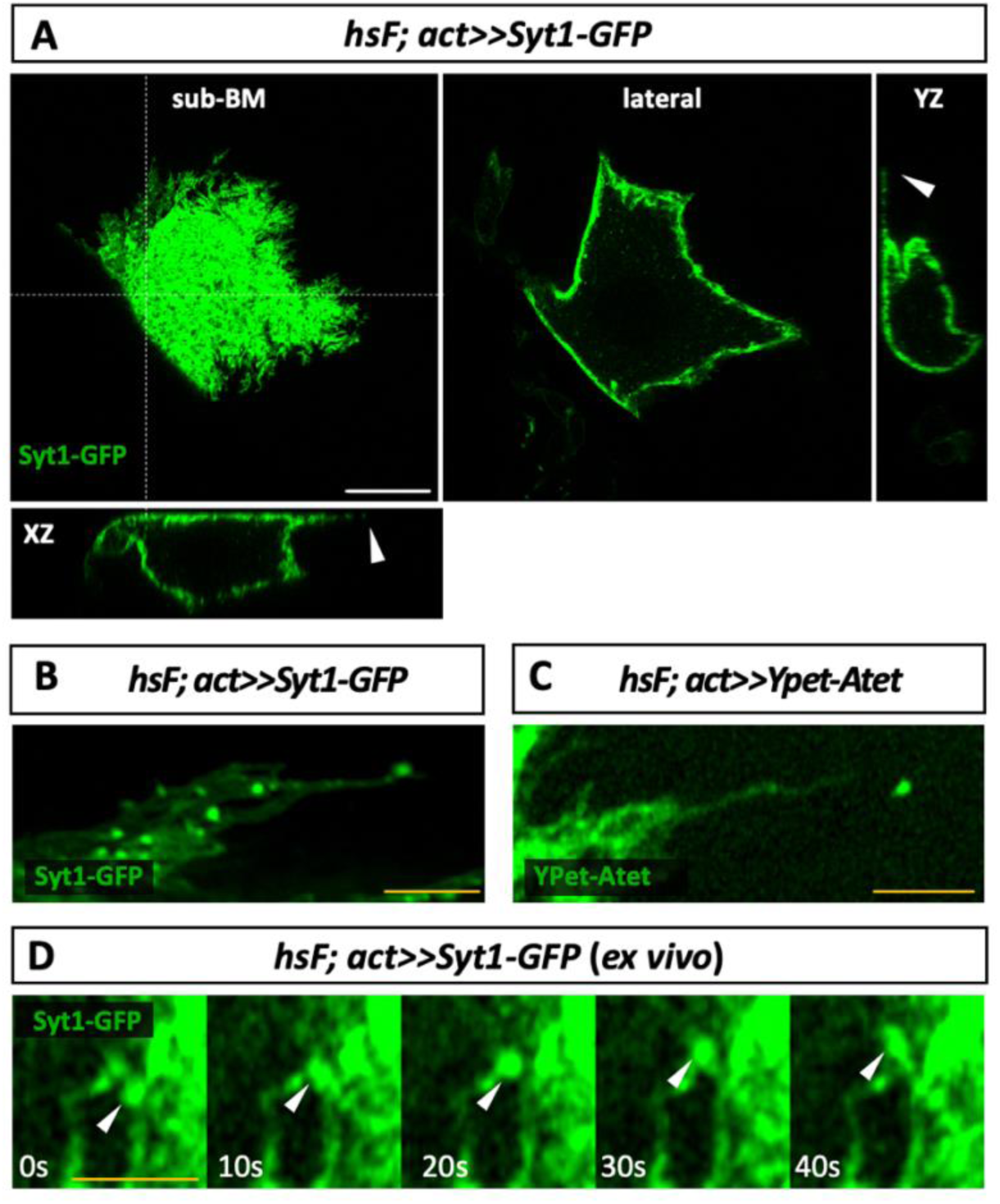
PG filopodia contains the machinery involved in Ecdysone secretion. **(A)** Syt1-GFP expressing clone showing Syt1 localized at the cell membrane and is enriched in filopodia in the sub-BM area. The arrows in the XZ and YZ sections point out filopodia illuminated by Syt1-GFP-expressing. **(B, C)** Zoomed-in images of filopodia displaying Syt1-GFP at the membrane and in vesicles were captured (B), as well as images showing Ypet-Atet at the membrane and in vesicles (C). **(D)** Tracking of a Syt1-GFP vesicle (arrow) in a filopodium. White scale bar: 20 μm; yellow scale bar: 5 μm.

### Filopodia mediate in Ecdysone secretion

To confirm the direct role of filopodia in Ecdysone secretion, we first disrupted filopodia, specifically in the PG, and analyzed the effects on developmental timing, which depends directly on secreted Ecdysone signaling. Using the PG-specific *phm-Gal4* driver, we conducted a bias RNAi screen targeting genes involved in filopodia structure. First, we manipulated the expression of α- and β-tubulin subunits that function as heterodimers. Misexpression of *αTub85E*, *βTub85D*, and *βTub56D* resulted in a larval arrest at the third larval stage or significant developmental delays of 35 and 50 hours, respectively (**Figure 5A**), indicating disrupted Ecdysone signaling. Interestingly, αTub85E is the α subunit shown in **Figure 2H** as specifically enriched in the PG cell cortex. Next, we destabilized the actin cytoskeleton (**Figure 5B**). Silencing actin (Act5C) with two independent RNAi constructs led to either L3 stage arrest or a 50-hour developmental delay. Disrupting actin polymerization or depolymerization by expressing chickadee (chic)^38^ or cofilin^29^ caused delays of 30 and 70 hours, respectively. Knocking-down Arpc3B, involved in actin branching^29^ and also explicitly enriched in ring gland^31^, resulted in L3 arrest. We confirmed the effects on filopodia morphology through αTub85E immunostaining (**Supp Figure 5**). Downregulation of actin (Act5C), or its polymerization (cofilin and chi) resulted in PGs with shortened filopodia, as shown by the visible cell edges. However, except for αTub85E, which shows specific localization at the cell cortex, altering the expression of the general cytoskeleton components such as act5C, cofilin, and chic might extend beyond filopodia, as these cytoskeleton proteins perform fundamental functions in all cells^21^.

**Figure 5.**
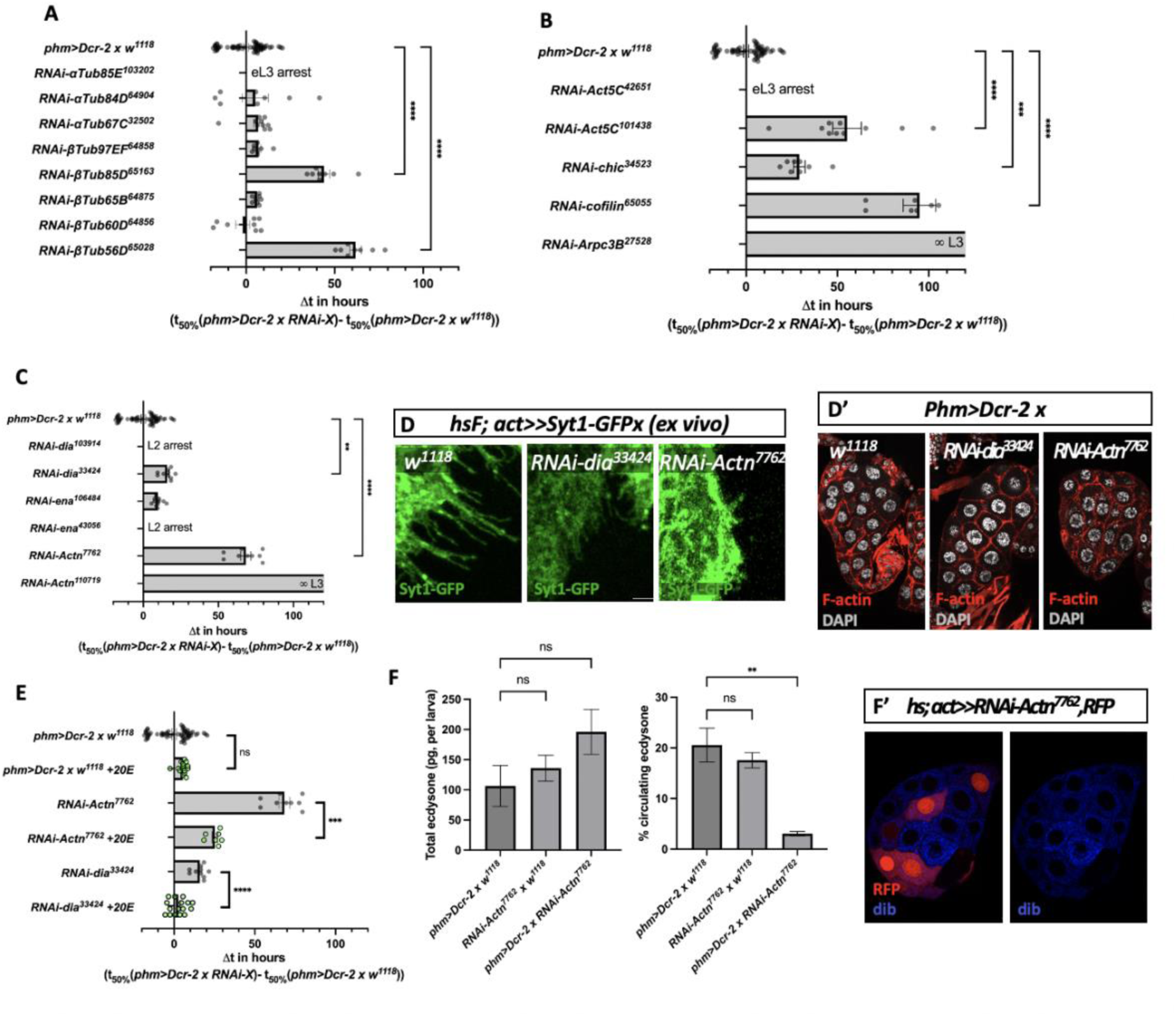
Alteration of PG filopodia components leads to Ecdysone secretion defects. **(A)** Screening for α- and βtubulin subunits involved in developmental timing by expressing the RNAi under the control of *phm-Gal4*. **(B)** Screening for general actin-associated proteins involved in developmental timing. **(C)** Screening for filopodia-associated proteins involved in developmental timing (**D,D’**). Compared to the control situation, inhibition of *dia* leads to shorter filopodia, while down-regulation of Actn leads to more severe filopodia defect (*ex vivo*, D). PG have an apparent normal morphology when dia or Actn is down-regulated (D’). **(E)** The developmental delay obtained upon inhibition of *dia* or *Actn* is rescued by feeding 20E, indicating the developmental delay observed is due to a lack of Ecdysone secretion. **(F,F’)** Circulating Ecdysone is strongly reduced when *Actn* is down-regulated compared to control (F). (F’) Dib expressing is not altered when *Actn-RNAi* is expressed. These results indicate that the removal of Actn does not influence Ecdysone production, but its secretion. Scale bar: 20 μm.

Therefore, we further altered the expression of cytoskeletal components already described as specific to filopodia (**Figure 5C**). Targeting the formin Diaphanous (dia) and Enabled (ena), both unbranched actin filament polymerases at the tips of filopodia ^20,39^, resulted in either L3 stage arrest or a 12-hour delay. Inhibition of α-actinin (Actn), an actin cross-linking protein at the base of filopodia^20,39^, also caused significant developmental timing disruptions, showing either L3 arrest or a 50-hour delay. Their effect on the filopodia network was confirmed by *ex vivo* experiments, which showed that when dia expression was down-regulated, there were shorter filopodia compared to the control group. A more significant effect on filopodia abrogation was observed when Actn was silenced in the PG (**Figure 5D; Supp Figure 5**). It is important to highlight that the PG maintains a regular shape and morphology (**Figure 5D’**). Interestingly, Ecdysone feeding of larvae with PG downregulated *dia* or *Actn* expression, our most significant filopodia-specific candidates, significantly rescued the previously observed developmental timing defects (**Figure 5E**). This indicates that the observed developmental defects were due to abnormal Ecdysone signaling. Furthermore, to determine whether this defect in Ecdysone signaling is due to a direct secretion defect, we determine Ecdysone secretory index as the percentage of Ecdysone present on circulating hemolymph relative to total larval Ecdysone in the wandering stage. In control conditions, around 20% of total Ecdysone circulates during the wandering stage (**Figure 5F**), as previously reported^40^. However, when *Actn* expression is downregulated in the PG, we observe high total ecdysone levels, showing that PG cells with reduced Actn produce Ecdysone efficiently; however, Ecdysone secretory index drops to only 3% of total larval Ecdysone (**Figure 5F**). We assessed disembodied (dib) expression, an ecdysone biosynthetic enzyme, in PG cells with random Actn-silenced clones to further confirm that Actn loss does not impair PG cell functionality. dib immunostaining showed similar expression levels between the wild type or upon Actn down-regulation (**Figure 5F’**). This confirms that specific PG silencing of Actn, a specific filopodia component, leads to Ecdysone secretion defects. These findings indicate that the filopodia sub-basal network is crucial in Ecdysone signaling, specifically in the cellular mechanisms governing Ecdysone secretion.

### Filopodia developmental dynamic correlates with the establishment of the large metamorphosis-inducing Ecdysone peak

Given the previous observation that several morphological changes occur in PG cells during late larval development ^13^, which coincides with a peak of Ecdysone production and secretion, we investigated whether filopodia exhibits developmental changes. We measured various filopodia parameters at early and late third instar larvae stages (eL3, lL3; **Figure 6A**) following the method analysis shown in **Supp Figure 6A**. We observed a significant increase in average filopodia length (**Figure 6B**) from approximately 1.8 μm in eL3 to 2.8 μm in lL3. In contrast, filopodia diameter showed no significant difference between the two stages (**Figure 6C**), ranging between 0.2 and 0.3 μm. Although the filopodia area increased between the early and late stages due to lengthening, each filopodia exchange surface relative to cell size remained similar between the two time points (**Supp Figure 6B**). Notably, we observed a substantial increase in filopodia density (**Figure 6D**) with 0.07 filopodia per μm^2^ in eL3 to 0.33 filopodia per μm2 in lL3. Taken together, filopodia at the sub-BM area surface increased the exchange surface by approximately 10% in eL3, whereas in lL3, the higher filopodia density led to a 70% increase in cell exchange surface (**Figure 6E**). Our results indicate that filopodia are necessary not only for secretion but also to increase the secretory surface in the sub-BM area of the PG just before a significant surge of Ecdysone is secreted into the hemolymph to induce pupariation.

**Figure 6.**
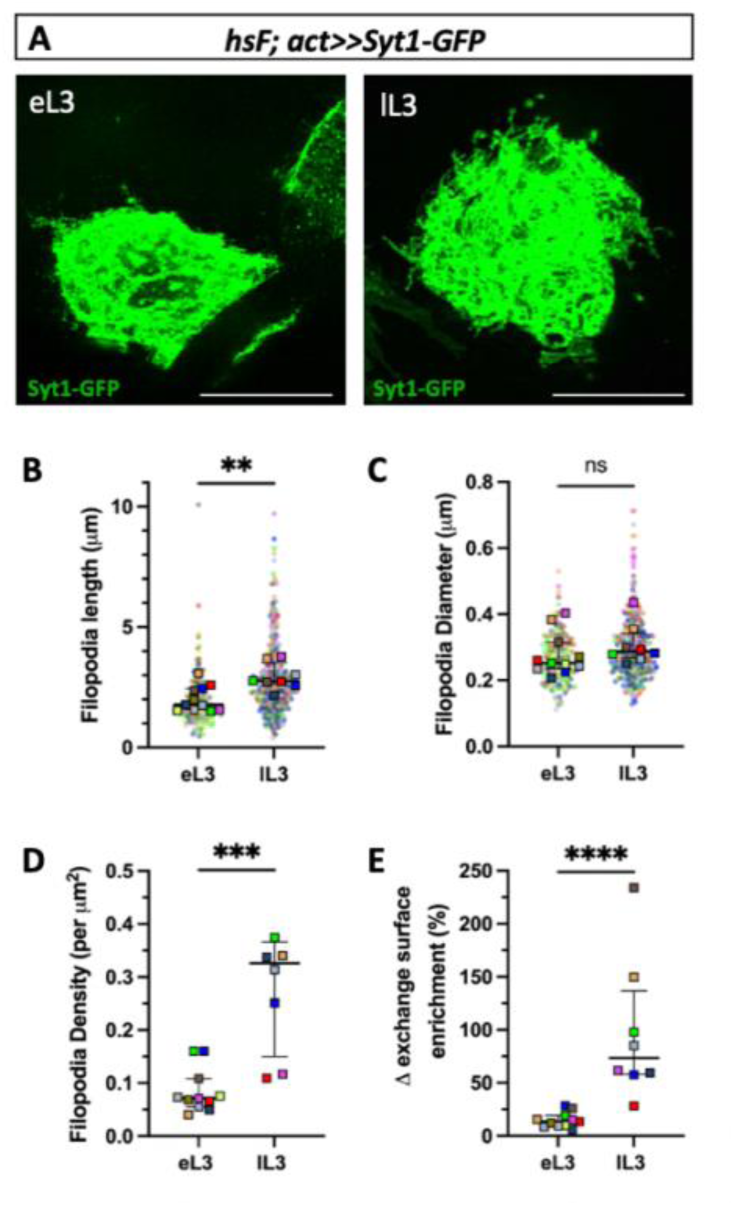
Filopodia increases the PG exchange surface at the sub-BM domain. **(A)** Representative images of an eL3 and lL3 PG cell expressing Syt-1-GFP. **(B-E)** Filopodia length increases significantly from eL3 to lL3 (B), while their diameter remains stable (C). The density of filopodia also increases significantly during the L3 stage (D), resulting in a 70% increase in the exchange surface of the sub-BM area (E). Scale bar: 20 μm.

## Discussion

### Hormonemes, filopodia-like structures: key players in ecdysone secretion

This work found that filopodia-like membrane projections, termed “hormonemes,” mediate Ecdysone secretion in the Drosophila prothoracic gland (PG). *ex vivo* experiments reveal that these structures contain vesicles marked by classical secretory proteins like Syt1 and Ecdysone-specific transporters such as Atet (**Figure 4B and C**). The enrichment of Syt-GFP at the filopodia structures (**Figure 4A**) and the significant drop in circulating Ecdysone levels following filopodia destabilization, despite intact Ecdysone production (**Figure 6F and F’**) confirm that these structures act as specialized secretory sites, forming a stable membrane network from which steroid-containing vesicles are released. Given the curved nature of filopodia-like structures, it is possible that they introduce localized membrane curvature that might optimize the energetic efficiency of vesicle-membrane fusion and thus enhance vesicle secretion.

### Polarized filopodia distribution as a mechanism for directed secretion

Our study reveals that the polarized distribution of filopodia serves as a novel mechanism for directed Ecdysone secretion outward the gland. PG cells lack distinct apical and basolateral domains, showing an almost uniform distribution of basolateral factors (**Figure 1E**). This absence of apico-basal polarity is also seen in other steroid-secreting glands, such as the zona glomerulosa rosettes of the adrenal cortex, which secrete aldosterone^41^. The lack of cell polarization in these highly secretory tissues raises an important question: How do these cells efficiently direct steroid hormones toward the circulatory system to elevate hormone levels rapidly? Our results indicate that the asymmetric distribution of filopodia on PG cells is essential for efficient hormone secretion. In PG cells, these filopodia are concentrated near the BM (**Figure 2B and C**). The directed movement of vesicles toward the filopodia tips (**Figure 4D**) suggests a directional secretion process. Indeed, one distinguishing feature we found on PG filopodia-like structures is their remarkable stability (**Movie 1**), attributed to interactions with the BM and tubulin within them (**Figure 2H and 3L**). This stability reduces their dynamics, forming a robust secretory platform near the gland’s outer layer. The identification of microtubules within PG filopodia suggests an essential role in polarized Ecdysone secretion. Supporting this, research in *Manduca sexta* has shown PTTH signaling increases β-tubulin synthesis in PG cells^42^ and suggests that microtubules should be important for steroid precursor uptake^43^. These findings imply that the asymmetric distribution of cytoskeletal components in PG cells could be vital not only for Ecdysone secretion but also for integrating local and systemic signals to control ecdysteroid biosynthesis, which requires further investigation.

### Evolutionary significance of filopodia-like projections

The results presented here raise the question about the evolutionary significance of these findings. When delving into the older literature, we found that the filopodia observed in our study resemble the “membrane invaginations” previously reported in the prothoracic gland (PG) cells of Drosophila melanogaster^44^, as well as similar structures in other insects like *Manduca sexta*^45^ and *Lymantria dispar*^46^. Electron microscopy studies showed these membrane invaginations as a network of membranes facing the BM, suggested to be involved in Ecdysone secretion^44^. However, earlier research struggled to define the membrane network accurately, proposing various interpretations like invaginations or channels possible due to the non-preservation of filopodia-like projections in the former studies compared with the current new protocols^47^. Our *ex vivo* or slightly fixed (see materials and methods) individual PG cell analysis protocol allowed us to identify filopodia-like projections. When observed in the context of all filopodia projections from all PG cells, they formed a membrane network similar to the one shown in the electron microscopy pictures. Moreover, these membrane structures close to the BM in *Lymantria dispar* carry secretory vesicles and are essential for Ecdysone secretion^46^, further supporting filopodia’s evolutionary conserved functional involvement in Insect steroid secretion.

### Could these filopodia-like structures be fundamental to hormone secretion across diverse organisms beyond insects?

This idea is supported by similar mechanisms observed in other glands: cholecystokinin (CCK) is secreted by neuroendocrine cells through pseudopod-like processes known as neuropods^48^, and L cells in the gastrointestinal tract extend neuropods to release peptide PYY^49^ ^50^. In insects, Inka cells, which secrete ecdysis-triggering hormones (ETH), harbor cytoplasmic extension^51^. In pancreatic delta cells, microtubule-containing filopodia facilitate somatostatin secretion^18^, and crustaceans have microvilli in the Y-organ that are suggested to be involved in the secretion of Ecdysone^50^. The parallels suggest that membrane extensions are universally used for efficient, targeted hormone secretion, highlighting their potential evolutionary significance. Our study enriches this understanding by emphasizing the role of filopodia in endocrine signaling within the Drosophila PG, providing new insights into the mechanisms of steroid hormone release.

## Acknowledgments

We thank the plantBios ISA microscopy facility for technical support. We thank G. de Soussa, G. D’Angelo, S. Pizette, and all BES laboratory members for insightful discussions and comments on the scientific work. We thank the Vienna Drosophila RNAi Center, the Drosophila Genetics Resource Center and the Bloomington Stock Center for providing Drosophila lines. We thank P. Leopold and F. Martin for the critical reading of the manuscript. We thank N.Yamanaka for providing us the *UAS-Ypet-Atet* and the *UAS-Dcr-2;phm22-Gal4* lines. We thank J.-C. Pastor-Pareja for providing us with the Cg25C antibody. We thank T. Perry for provinding us the *Phm-Gal4* line. This work was supported by the ATIP_AVENIR young group leader program (INSERM grant 2020/5 to N.M.R), the Marie Curie fellowship (EU MSCA to E.S), and the Labex Signalife program (grant NR-11-LABX-0028-01 and IDEX UCA Jedi ANR-15-IDEX-01 to N.M.R.).

## Author contributions

E.S., R.B., Y.M., and M.-P.N.-E. conducted the experiments. E.S., R.B., and N.M.R. designed and interpreted them. E.S. wrote the original manuscript, and E.S., R.B., and N.M.R. wrote, edited, and reviewed it. N.M.R supervised the work and led the conceptualization. All the authors were involved in the scientific discussions.

## Material & Methods

### Drosophila strains and husbandry

Flies were reared and crossed at 25°C on standard cornmeal food containing, per liter, 15g inactivated yeast powder, 80g corn flour, 8g agar (Sigma, A7002) and 3,5g Nipagin (Sigma, W271004). Recombinant or combination stocks were made using standard genetic methods. The following fly strains were used: *w^1118^; UAS-Dcr-2; phm22-Gal4* (Gift of Naoki Yamanaka), *w^11118^; phm-Gal4*^52^, *P0206-Gal4*^53^, *UAS-myr-RFP* (BDSC7119), *UAS-myr-GFP* (BDSC32197), *UAS-PLCδ-PH-EGFP* (BDSC39693), *hsF; UAS-PLCδ-PH-EGFP* (Gift of Laura Boulan), *UAS-syt1-GFP* (BDSC6925), *UAS-sec-GFP*^54^, *CoinFLP LexGAD/Gal4,LexAop-CD2-RFP; UAS-CD4-spGFP^1-10^,LexAop-CD4-spGFP^11^* (BDSC58755), *hsFLP; UAS-CD8-RFP* (BDSC27392), *Trol-GFP*^55^, *vkg-GFP* (DGRC110692), *Ndg-GFP* (VDRC318629), *LanB1-GFP* (VDRC318180), *baz-GFP* (BDSC51572), *crb-GFP* (BDSC99495), *Sgs3-GFP* (BDSC5884), *UAS-RNAi-cofilin* (BDSC65055), *trol^JO271#49^* ^56^, *UAS-RNAi-trol* (BDSC38298), *UAS-RNAi-rab11*, *UAS-RNAi-Vps25*, *UAS-RNAi-Vps36*, *UAS-RNAi-shrub*, *UAS-RNAi-mys*, *Ypet-Atet*^57^, *UAS-RNAi-dia* (103914KK and BDSC33424), *UAS-RNAi-ena* (106484KK and BDSC43056), *UAS-RNAi-Actn* (7762GD and 110719KK), *UAS-RNAi-Act5C* (BDSC42651 and 101438KK), *UAS-RNAi-chic* (BDSC34523), *UAS-RNAi-Arpc3B* (BDSC27582), *UAS-RNAi-αTub85E* (103202KK), *UAS-RNAi-αTub84D* (BDSC64904), *UAS-RNAi-αTub67C* (BDSC32502), *UAS-RNAi-βTub97EF* (BDSC64858), *UAS-RNAi-βTub85D* (BDSC65163), *UAS-RNAi-βTub65B* (BDSC64875), *UAS-RNAi-βTub60D* (BDSC64856), *UAS-RNAi-βTub56D* (BDSC65028).

### Developmental timing experiments

All developmental timing experiments were conducted at 25°C. Embryos were collected after 4 hours of egg laying carried out on agar (1,5% w/v) dishes containing 3% (w/v) sucrose and were immediately transferred into tubes containing standard cornmeal food at a rate of 50 individuals by tube. The percentage of pupation in function of time was determined for each tube by counting pupae number two times a day.

For the rescue by 20-HydroxyEcdysone feeding, 50 μL of a freshly prepared solution (0,25mg/mL in ethanol absolute, stock solution at 5mg/mL, sigma H5142) were directly added on the top of the food twice a day, starting at late second instar stage (before 72hAEL).

### Fluorescent immunostaining

Third instar larvae were dissected in PBS, fixed in 4% formaldehyde (16% stock solution, PolySciences 18814-20) in PBS for 25mn at R.T., permeabilized with 0,1% Triton X-100 PBS three times and blocked in 1X ROTI^®^-Block (Stock solution 10X, Carl Roth A151.1) 0,1% Triton X-100 PBS for 40mn. Dissected larvae were incubated in blocking solution containing primary antibodies overnight at 4°C, washed three times in blocking solution and incubated with fluorescent secondary antibodies diluted in blocking solution for 2h at R.T. Samples were washed three times in PBS and mounted in Vectashield mounting media with DAPI (Vector Laboratories H-1200).

Primary antibodies were diluted as follows: guinea pig monoclonal α-PTTH (1:400)^58^, rabbit polyclonal α-Torso (1:200, this study), guinea pig polyclonal a-disembodied (1:400, this study), mouse monoclonal α-Arm (1:50, DSHB N27A1), rat monoclonal α-Ecad (1:50, DSHB DCAD2), α-Cora (1:50, DSHB C615.16), mouse monoclonal α-Dlg1 (1:50, DSHB 4F3), mouse monoclonal α-βPS (1:50, DSHB CF.6G11), rat monoclonal α-Ncad (1:50, DSHB DN-Ex #8), rat polyclonal α-Crb2.8 (1:300)^59^, mouse monoclonal α-αTub85E (1:50, DSHB 12G10), rabbit polyclonal α-Trol (1:3500)^60^, rabbit polyclonal α-Cg25c (1:1000)^61^. Polyclonal antibodies against Torso were raised using the following peptide for rabbit immunization (Eurogentec): -WVQHSRGTEPAPNAT- and -PKRKLKPQPKKDSKQ-. Polyclonal antibodies against disembodied (dib) were raised using the following peptide for guinea pig immunization (Eurogentec): -CKTLLINKPDAPVLIDLRLRRE-. Fluorescent secondary antibodies (Alexa Fluor or Fluor plus 488-, 546- or 555-, 647-conjugated antibodies from ThermoFisher Scientific) and TRITC-conjugated phalloidin (Sigma P1951) were diluted at 1:200. To compare specimens of different genotypes, samples were stained in parallel using the same antibody solutions or in the same tube when tissue morphology allowed them to be distinguished, and imaged with identical acquisition settings. Confocal images were acquired on a Zeiss LSM 880 with Airyscan using the Fast mode of the Airyscan detector. X.Y. pictures are either single optical sections or Maximum Intensity Projections obtained from a z-stack acquisition. XZ and YZ pictures are orthogonal sections extracted from a z-stack acquisition. All images were processed and analyzed with Fiji software. 3D visualization of the ring gland was obtained with Vaa3D software ^62^.

### PG cells number

PG cells were numbered by manually counting nuclei stained with DAPI. Belonging to the dorsal or ventral layer of the PG was determined using z-stack acquisition.

### *ex vivo* imaging and live imaging

The anterior part of late third instar larvae was dissected and directly mounted between two coverslips of different sizes in HL-3 saline solution (70 mM NaCl, 5 mM KCl, 1.5 mM CaCl2,20 mM MgCl2, 10 mM NaHCO3, 5 mM trehalose, 115 mM sucrose and 5 mM HEPES, pH 7.4). Confocal images were acquired on a Zeiss LSM 880 with Airyscan using the Fast mode of the Airyscan detector. Live imaging was achieved using a time series scan, with an interval of 10s between 2 frames.

### Ecdysone titration

#### Isolation of exosomes from hemolymph

This protocol was adapted from^32^. Briefly, 20μL of hemolymph were collected from late third instar larvae, transferred in 500µl of PBS+PTU (phenylthiourea 0,1µg/ml, Merck, P7629), and centrifuged at 750g for 15min at 4°C. The supernatant was transferred into a new tube and centrifuged again at 1500g for 10 minutes at 4°C. The remaining supernatant was then centrifuged a last time at 100 000g for 2h at 4°C. The final pellet was suspended in 300µl of methanol for Ecdysone titration. Methanol samples were centrifuged at 12 000g for 5min at 4°C. First, the supernatant was kept, and the pellet was suspended again in 300µl of methanol and centrifuged in the same conditions. The second supernatant was pooled with the first and stored at −80°C.

#### Ecdysone extraction from hemolymph and total larva

For hemolymph Ecdysone, 5 to 7L of hemolymph were collected from late thirst instar larvae and resuspended in 90µl of PBS+PTU. For total larva Ecdysone, four late third instar larvae were disrupted in 100µl of PBS+PTU with a Tissue Lyser (Qiagen) at 50Hz for 1min and centrifuged at 2000g for 10min at 4°C, and the supernatant was kept.

300µl of methanol were added to both samples and centrifuged at 12 000g for 5 min at 4°C. The first supernatants were kept, and pellets were suspended again in 300µl of methanol and centrifuged in the same conditions. The second supernatants were pooled with the first and stored at −80°C.

#### Ecdysone titration assay

Methanol from isolated exosomes, hemolymph, or total larva samples was removed using a Speed Vacuum for 2,5h. Dried pellets were resuspended in 120µl of EIA buffer from the 20-HydroxyEcdysone ELISA kit (Bertin Pharma, A05120). Ecdysone titration was then performed according to the manufacturer’s protocol.

### Filopodia measurements

Synchronized larvae of the indicated genotype (*hsF; act>>syt1-GFP* or *hsF; act>>PLC-GFP*) were heat-shocked at 37°C for 10mn and allowed to rest at 25°C for 24h prior to dissection. Early or late third instar larvae were dissected in PBS and fixed in 4% formaldehyde for 10mn at R.T. After three washes in PBS, PG were mounted in Vectashield mounting media with DAPI and assessed for the presence of isolated syt1-GFP or PLC-GFP expressing clones, to properly visualize the filopodia. Confocal images of isolated PG clones were acquired on a Zeiss LSM 880 with Airyscan using the Fast mode of the Airyscan detector. Maximum Intensity Projections of the slices corresponding to the sub-BM region were manually analyzed on ImageJ software. Sub-BM cell perimeters and areas without filopodia were quantified and, thanks to the contrast with the surrounding “black-background”, filopodia located at the edge of the clone were counted and measured in their length (L) and diameter (d).

The number of filopodia was normalized to the perimeter on which they have been counted to assess filopodia density per μm and this value was used to estimate a filopodia density per μm^2^. Assuming the distribution of filopodia was homogenous throughout the entire sub-BM surface, the number of filopodia for a given clone cell was calculated by multiplying filopodia density (per μm^2^) by cell sub-BM area.

The exchange surface of each counted filopodia was calculated using the formula for the total surface area of a cylinder from which area of one circular base have been subtracted:

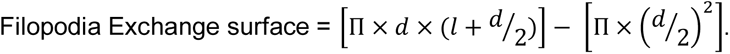

This filopodia exchange surface was also normalized to the cell sub-BM area and this parameter is referred to as “Filopodia exchange surface (relative to cell size)”

For a given clone cell, the Δ exchange surface enrichment was calculated using the following formula:

Δ exchange surface enrichment (in %) = (Exchange surface with filopodia – Exchange surface without filopodia) x 100,

in which:

Exchange surface with filopodia = (number of filopodia x average exchange surface of 1 filopodia) + cell sub-BM area;

Exchange surface without filopodia = cell sub-BM area.

### Fluorescence intensity measurement

Confocal images of syt1-GFP clone cells were obtained using the same protocol than the one described above (see “Filopodia measurements” section). At least six distinct orthogonal sections (three in the XZ plane and three in the YZ plane) of a clone were used to determine syt1-GFP intensity at three different cell regions (referred to as sub-BM, lateral, and midline). Measures were done within the ROI box represented in Figure 4H, and the six values were averaged to obtain syt1-GFP intensity at a given region.

### Statistical analysis

All statistics and plots were obtained with the GraphPad Prism software. For Figure 1F, datasets are represented in a paired aligned dot plot displaying the mean ± 95%CI. Each dot represents fluorescence intensity for one sample, and the pairing line between dots represents the belonging to the same sample. In Figures 3A, B, N and S3F, developmental timings of the different genotypes were displayed in an X.Y. graph in which means ± SEM of the proportion (% pupariation) of all the tubes were plotted in function of time (in hours) after egglaying (abbreviated hAEL). In Figures 5A, B, C and E the differences of time at 50% of pupariation (abbreviated t_50%_) between the different experimental conditions (*phm>Dcr-2,RNAi-X)* and the control (*phm>Dcr-2/w^1118^)* were compared in a bar plot displaying the mean of the differences ± SEM. Each dot represents the value for one tube. For Figures 5F, S1A, and S3A, datasets are also represented by bar plots showing the mean ± SEM. In Fig S1A, each PG is color-coded, and each colored dot represents the cell number for one given PG. In Figure S3A, each dot represents the value for one sample. For Figures 6B, C and S6B, datasets are represented by superplots in which each clone cell is color-coded. The colored dot represents individual filopodia for a given clone cell, and the corresponding colored-square are meant for the different clone cells. The median of the means and interquartile range (IQR) are displayed in those plots. For Figures 6D and E datasets are represented by scatter dot plots indicating the median and the IQR. Each sample is color-coded, and each color-coded square represents the average for a given clone cell. For Figure 1F, data were subjected to the Friedman test followed by Dunn’s multiple comparison tests. For Figure S1A, data were subjected to the matched-pairs Wilcoxon test. For Figures 5A, B, C and 5F, data were subjected to the Kruskal-Wallis test, followed by Dunn’s multiple comparison tests. For Figures 5E, 6B, C, D, E and S6B, data were subjected to the Mann-Whitney test. The significance of statistical tests is reported in plots as follows: NS: not significant, *: p < 0.05, **: p < 0.01, ***: p < 0.001, ****: p < 0.0001.

