## Supplemental data for "Filopodia-like Structures are Essential for Steroid Release"

**Supplementary figures**

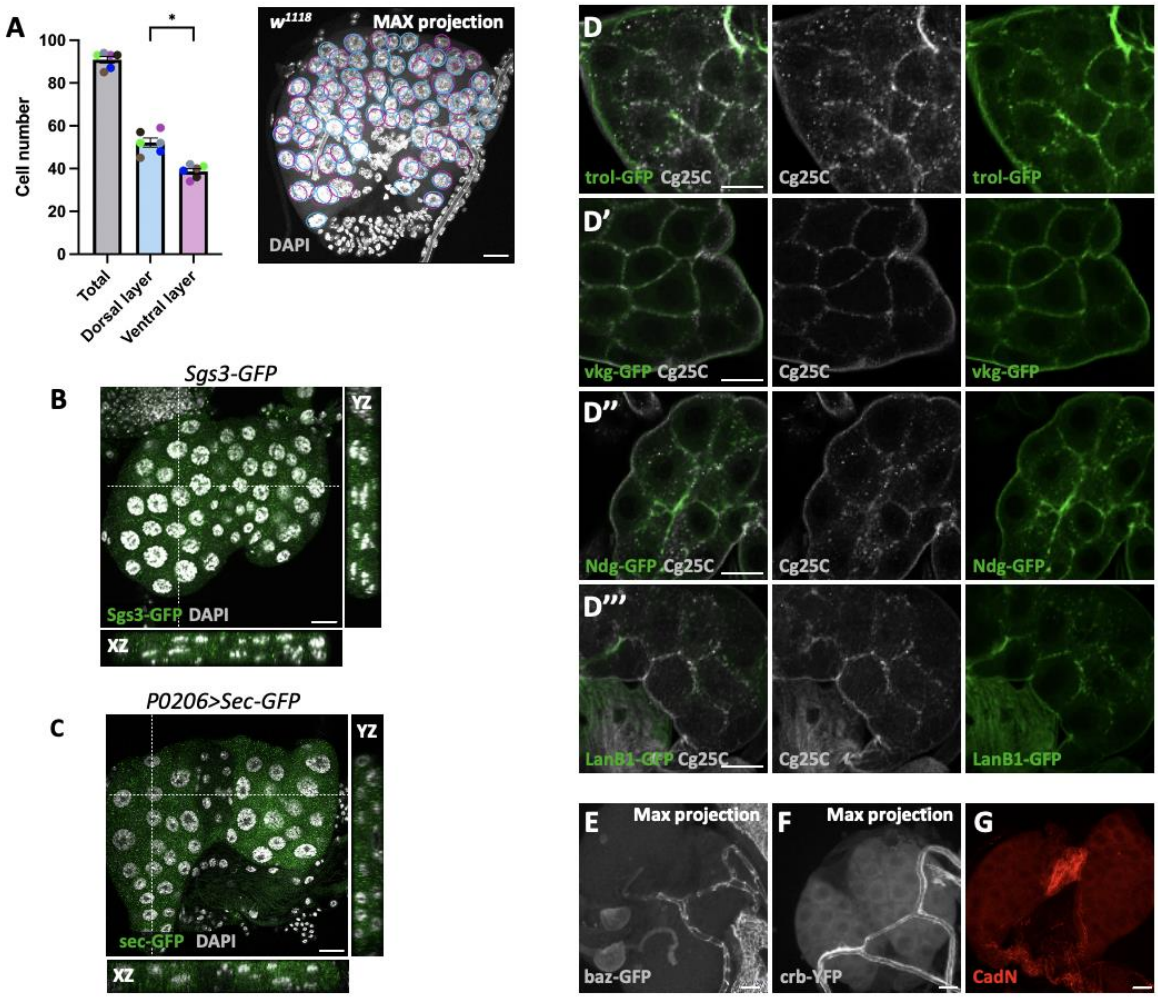


**Supp Figure 1. PG general organization.** (**A**) Number of PG cells in each layer (n=6). (**B, C**) Sgs3-GFP and sec-GFP do not accumulate at the PG midline, indicating that PG is devoid of lumen. (**D-D'''**) trol-GFP (D'), vkg-GFP (D''), Ndg-GFP (D''') and LanB1 (D'''') colocalized with Cg25C at the BM. A colocalization is also observed in dots in the extracellular space indicating those structures are CIVICs. (**E, F**). The apical markers baz-GFP (E) and crb-YFP (F) are not expressed in the PG cells; their expression is only detected in the trachea. (**G**) CadN is not expressed in the PG but only in the CA Scale bar: 20 µm.

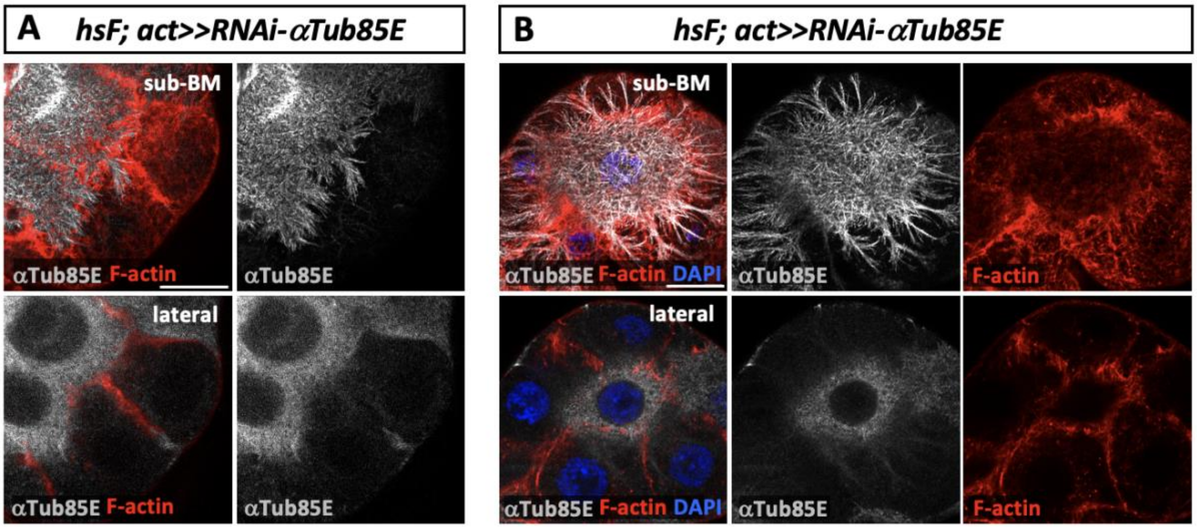


**Supp Figure 2. PG filopodia contain tubulin.** (**A**) Staining of αTub85E in a PG having few RNAi-αTub85E clones. Microtubules from WT cells are revealed when the neighboring mutant cells lack αTub85E. (**B**) A PG showing a single WT cell surrounded by RNAi-αTub85E clones. This WT cell shows longer microtubules, indicating a possible plasticity of these filopodia. Scale bar: 20 µm.

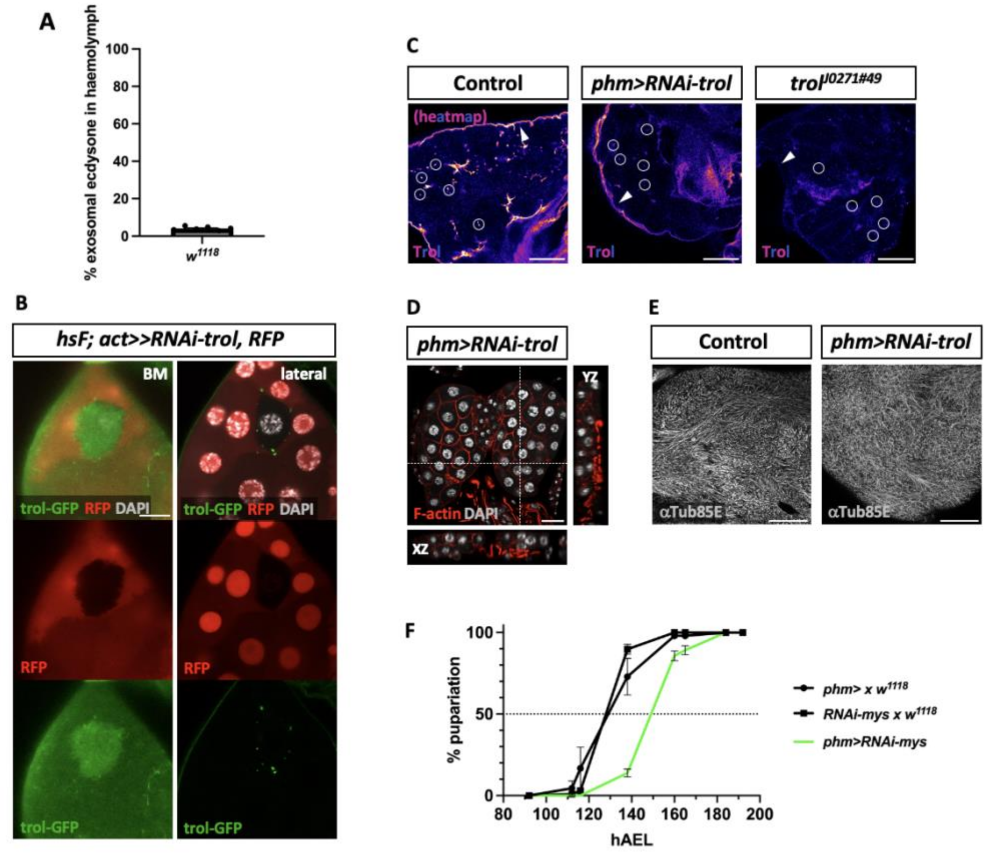


**Supp Figure 3. Analysis of exosomal Ecdysone and autonomous and systemic *trol* contribution on BM and CIVIC PG structures.** (**A**) Quantification of circulating Ecdysone indicates that less than 5% is found in the exosomal fraction. (**B**) Flip-out clones expressing an RNAi against *trol* and being positively marked RFP. A single WT cell (RFP negative) surrounded by mutant *RNAi-trol*-expressing cells is observed. On the BM plan, a decrease of *trol-GFP* is observed when the *RNAi-trol* is expressed. In the lateral plan, a complete lack of CIVICs is seen in the mutant cells. CIVICs are only present surrounding the WT cell. These cells belong to the dorsal layer of a PG. (**C**) Highlighting *trol* expression in the BM and CIVICs with a heatmap color code in a control situation (left picture). When *trol* expression is down-regulating specifically in the whole PG, *trol* expression is still observed in the BM while its expression strongly decreases in the CIVICs (middle picture). In a *trol* hypomorphic mutant, *trol* expression is barely visible in the BM as in the CIVICS (right picture). The arrows point at the BM, and the circles indicate the CIVICs. (**D**) Down-regulation of *trol* expression, specifically in the PG, does not alter the PG bilayer organization. (**E**) Control PG showing microtubules in the sub-BM region. A similar pattern is observed when *trol* is down-regulated specifically in the PG. (**F**) Developmental timing of *phm>RNAi-mys* vs controls. Down-regulation of βPS lead to a 24 hours developmental delay. Scale bar: 20 μm.

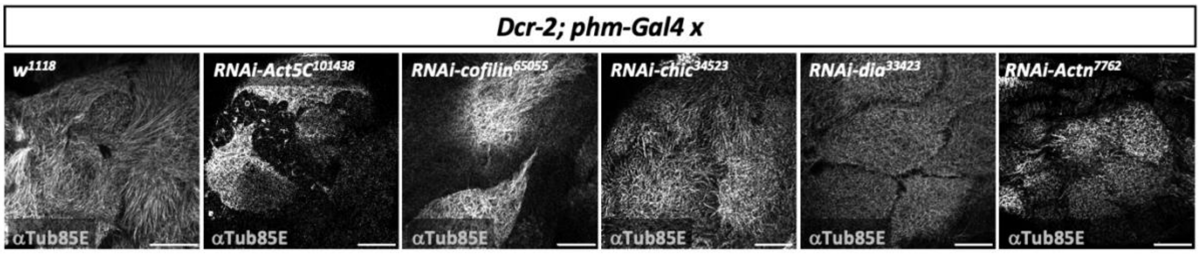


**Supp Figure 5. The microtubule network is altered when the actin machinery is disturbed.** Immunostaining of α-Tub85E in PG expressing different RNAi, which have shown developmental delay, alters the actin cytoskeleton. Scale bar: 20 µm.

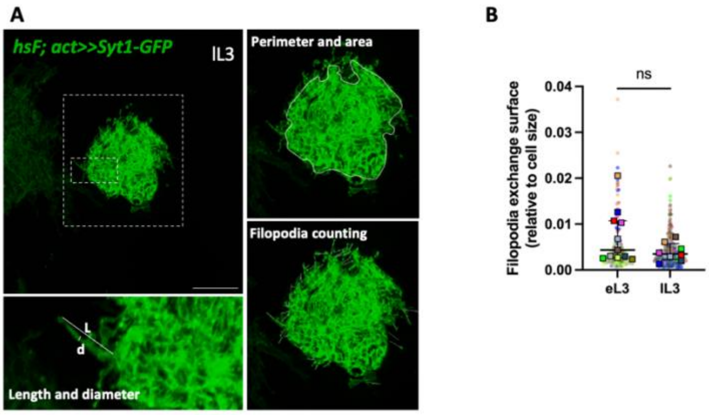


**Supp Figure 6. Quantification of filopodia exchange surface through development.** (**A**) The methodology used for filopodia quantification analysis. Different filopodia parameters were measured at two time points: early L3 (eL3) and late L3 (lL3) using the Syt1-GFP marker (**B**) Relative to cell size, the filopodia exchange surface is stable over time.
